# Neural sensitivity to radial frequency patterns in the visual cortex of developing macaques

**DOI:** 10.1101/2025.01.27.634810

**Authors:** C. L. Rodríguez Deliz, Gerick M. Lee, Brittany N. Bushnell, Najib J. Majaj, J. Anthony Movshon, Lynne Kiorpes

**Author notes:** **Correspondence:** Lynne Kiorpes, Center for Neural Science, New York University, 4 Washington Place Room 621, New York, NY 10003.

## Abstract

Visual resolution, contrast sensitivity and form perception improve gradually with age. In nonhuman primates, the sensitivity and resolution of cells in the retina, lateral geniculate nucleus and primary visual cortex (V1) also improve, but not enough to account for the perceptual changes. So, what aspects of visual system development limit visual sensitivity in infants? Improvements in behavioral sensitivity might arise from maturation of regions downstream of V1 such as V2, V4 and pIT, which are thought to support increasingly complex perceptual abilities. We recorded the responses of populations of neurons in areas V1, V2, V4, and pIT to radial frequency patterns - a type of global form stimulus. Subjects were three young monkeys between the ages of 19 and 54 weeks, and a single adult animal. We found that neurons and neural populations in V4 reliably encoded global form in radial frequency stimuli at the earliest ages we studied, while V1 neurons do not. V2 and pIT populations also showed some degree of selectivity for these patterns at early ages, especially at higher radial frequency values. We did not find significant, systematic changes in neural decoding performance that could account for the improvement in behavioral performance over the same age range in an overlapping group of animals (Rodriguez Deliz et al., 2024). Finally, consistent with our prior behavioral results, neural populations in V4 show highest sensitivity for the higher radial frequency values which contain the highest concentration of curvature and orientation cues.

**SIGNIFICANCE STATEMENT:** Infants have remarkably limited ability to discriminate shapes. These limitations cannot be fully explained by postnatal changes in their eyes, visual thalamus, or primary visual cortex. The perception of shape requires integration of local cues across space to create global form information. We therefore examined populations of neurons in extrastriate visual cortex to learn whether information represented in these regions might limit infants’ abilities to process global forms. We found instead that extrastriate areas involved in global form processing function maturely early in life, by the age of 4-6 months, suggesting that infants’ perceptual limits are set by other aspects of brain development.

## INTRODUCTION

Many features of visual perception, including spatial resolution, contrast sensitivity, and global motion and form perception, are poor in early infancy and improve gradually with age (Kiorpes, 2016; Kiorpes and Movshon, 2004; Rodriguez Deliz et al., 2024). In non-human primates, this improvement cannot be fully explained by the functional maturation of cells in the retina, lateral geniculate nucleus, or primary visual cortex (V1) (Danka Mohammed and Khalil, 2020; Kiorpes, 2016; Kiorpes and Movshon, 2004). The mechanisms that limit infant visual sensitivity therefore remain unclear.

Object perception involves hierarchical processing stages. The visual system extracts increasingly complex stimulus features from local cues such as oriented edges, angles and curves and integrates them across multiple spatial scales to create global form percepts (Kimchi, 1992; Kimchi et al., 2005; Van Essen et al., 1992). One possible explanation for perceptual improvements throughout development could be ongoing maturational cascades throughout the extrastriate cortex (Danka Mohammed and Khalil, 2020; Guillery, 2005; Kiorpes and Movshon, 2004).

Some understanding of development along the visual processing streams beyond V1 comes from studies of human infants using non-invasive electrophysiological methods. Visually-evoked potential recordings (VEPs) in children show weaker responses to global form stimuli compared to global motion at 8-10 weeks (Braddick and Atkinson, 2007). High-density VEP mapping confirmed these findings and revealed spatially distinct voltage distributions for form and motion stimuli in both infants and adults (Wattam-Bell et al., 2010). Evidence suggests that global motion processing (dorsal stream) emerges earlier but develops more slowly than global form processing (ventral stream) (Braddick and Atkinson, 2011; Braddick et al., 2003). These findings are consistent with behavioral data from nonhuman primates showing a similar pattern of relative development of sensitivity to global motion and form (Kiorpes et al., 2012) and anatomical data on differential development of dorsal and ventral streams (Distler et al., 1996).

Understanding how the visual cortex transforms and integrates stimulus information into reliable representations requires us to explore the computations carried out by visual neurons in more detail. Macaque monkeys have visual systems similar to that of humans, allowing us to track stimulus-evoked activity both throughout the visual system and across development. In macaques, VEPs to concentric and radially-organized Glass patterns emerge later (20-30 weeks) than texture-related VEPs (10 weeks) (Voyles, 2015). Physiological evidence from adult macaques reveals specialized neurons in areas V2, V4, and IT that encode increasingly complex stimulus features (Desimone et al., 1984; Desimone and Schein, 1987; Freeman et al., 2013; Gross and Rodman, 1992; Hegdé and Van Essen, 2000, 2003, 2007; Lee et al., 2024; Maruko et al., 2008; Okazawa et al., 2017; Rodman et al., 1993; Tanaka, 1996; Van den Bergh et al., 2010; Van Essen and Gallant, 1994; Ziemba et al., 2016, 2024).

The mid-level cortical region V4 may play a key role in global form processing (El-Shamayleh and Pasupathy, 2016; Gustavsen and Gallant, 2003; Merigan, 1996; Nandy et al., 2013; Pasupathy, 2006; Pasupathy and Connor, 1999, 2002; Roe et al., 2012; Van Essen et al., 1990; Young, 1992). Single-cell studies in monkeys have confirmed V4 cells’ responsiveness to concentric structures and contour features of varying concavity and convexity amplitudes (Gallant et al., 1996; Kim et al., 2019; Kobatake and Tanaka, 1994; Pasupathy, 2006; Pasupathy and Connor, 1999, 2002; Popovkina et al., 2019). Many V4 cells display size-independent sensitivity to form, with stimulus size possibly modulating response gain (El-Shamayleh and Pasupathy, 2016). V4 neurons also exhibit varying degrees of sensitivity to local polar phase cues (Nandy et al., 2013). V4 has larger receptive fields than earlier visual areas, allowing for integration of local elements across visual space (Felleman and Van Essen, 1991; Gattass et al., 1988; Maunsell and van Essen, 1983; Wilson and Wilkinson, 2004). Beyond V4 lies area IT, which responds to complex stimulus features like objects, faces, and color-object combinations (Desimone et al., 1984; Gross, 1992, 2008a,b; Gross et al., 1972; Gross and Rodman, 1992; Hung et al., 2012; Perrett et al., 1982; Tanaka et al., 1991; Tsao and Livingstone, 2008). Both V4 and IT neurons carry more information about natural images than their texturized counterparts, suggesting that they may encode the global form structure present in visual scenes (Rust and DiCarlo, 2010).

Here we explore the encoding of global form stimuli at multiple levels of the extrastriate pathway during development in young monkeys, in comparison to improvements in psychophysical performance as documented in another study (Rodriguez Deliz et al., 2024). We conducted longitudinal multi-electrode recordings in areas V1, V2, V4, and pIT of awake, fixating macaques. We used radial frequency stimuli (RFS) (Wilkinson et al., 1998), the perception of which requires global processing mechanisms (Bell et al., 2007; Jazayeri and Movshon, 2006; Poirier and Wilson, 2006; Tolhurst et al., 1983). Previous work in human and macaque infants has shown that sensitivity to these patterns is poor at young ages and improves over an extended time course (Birch et al., 2000; Rodriguez Deliz et al., 2024), suggesting that they represent a good metric for characterizing the neural mechanisms underlying the development of global form perception. We expected young V1 to be primarily sensitive to local orientation elements, while V4 and pIT cells would show sensitivity to global stimulus features. To test whether developmental changes in V4 or beyond account for behavioral improvements observed in (Rodriguez Deliz et al., 2024), we used population decoding methods to determine if decoding accuracy improved significantly with age. Our findings suggest that the limiting factor on infant global form perception may lie elsewhere in the cortex.

## MATERIALS AND METHODS

### Subjects

We studied 3 infant and 1 adult, visually normal pig-tailed macaque monkeys (*Macaca nemestrina*), 2 female and 2 male, between the ages of 19 and 398 weeks (5 months to 7.6 years old). Three infant subjects were recorded multiple times between the ages of 19 and 52 weeks; the adult was recorded at one age. Subject details are presented in Table 1. All animal care, surgical and experimental procedures were approved by the Institutional Animal Care and Use Committee of New York University and were in compliance with the guidelines established in the United States National Institutes of Health Guide for the Care and Use of Laboratory Animals.

**Table 1:** Subjects. Identification, cortical areas recorded from and range of age points included in this study are reported per subject. Since delimitations of the foveal confluence were not always clear at the time of implantation, recordings from M3 may reflect neural activity in V1, V2 or V4. Recording ages are reported for time points at which at least one array yielded reliable data. Subject M17 was tested monocularly with a +1.0D lens.

| Subject | Areas recorded | Recording ages (wk) |
| --- | --- | --- |
| M1 (female) | V1/V2 border, V4 | 31, 32, 38, 48, 54 |
| M2 (female) | V2, V4 | 29, 36, 46, 53, 54 |
| M3 (male) | Foveal confluence, pIT | 19, 20, 21, 22, 23, 24, 26, 32 |
| M17 (male) | V1, V4 | 398 |

### Neural recordings

We used platinum-tipped multi-electrode “Utah” recording arrays with 96 microelectrodes in a 10×10 grid pattern (excluding corners), separated by 400µm on a 3.6 × 3.6mm surface (Blackrock Microsystems, Salt Lake City, UT). The target sites for the arrays on the cortical surface were located using sulcal (lunate, inferior occipital and superior temporal sulci) landmarks during surgery to ensure implantation of the array in cortical regions representing the center of gaze. Target locations (see Results, Figure 2) for the V1/V2 array were directly behind the lunate, using the clear change in vasculature patterns at the border between V1 and V2 as a reference (Zheng et al., 1991). Target V4 and pIT locations were between the lunate and the superior temporal sulcus, toward the tip of the inferior occipital sulcus.

Our experimental plan involved collecting physiological data as early as possible for the 3 infants, which precluded waiting the required time for osseointegration of a headpost. The infants were therefore not head-fixed but were tested using custom side panels to restrict head movement; the adult was fitted with a headpost. Data collection began after a 2–3-week post-array-implantation recovery period. Data collection continued as long as array recording quality was satisfactory.

Neural signals were recorded continuously throughout each session using Blackrock CerePlex E headstages at a sampling rate of 30 kHz using Neural Signal Processors (Blackrock Microsystems). The impedance of the microelectrodes ranged from 200-800 kΩ. Signals were amplified and filtered in real time using a second order digital Butterworth filter (50 Hz - 250 kHz) and this band was digitized into ‘spikes’ based on threshold multi-unit events. Data were logged using Blackrock Central software (Blackrock Microsystems, Salt Lake City, UT). Thresholds were set at the beginning of each recording session by re-referencing data to the median voltage across all channels on the array. Threshold crossing events were defined as points when a channel’s voltage crossed -3.5x the RMS of its baseline activity. We extracted thresholds using a range of standard deviations (3-5) for a subset of our data and found that the threshold criterion choice did not influence our findings. For our purposes, threshold crossings and firing rates are used interchangeably. Firing rates described below reflect the activity of ensembles of neurons near the tip of the electrode, not individual cells. The first several recording sessions consisted of testing the array quality by assessing the proportion of active channels on each array. Baseline responsiveness of each channel was measured based on its response to a blank gray screen (as detailed below).

### Stimuli

Radial frequency stimuli (RFS), illustrated in Figure 1, are a family of closed contours created from a circle whose radius is sinusoidally modulated at different frequencies and amplitudes (see Figure 1B). The amplitude of these modulations can be parametrically adjusted to change the similarity between the stimulus and a base circle. In our case, the cross-section of the outline’s luminance profile was a narrowband stimulus defined by the fourth-derivative of a Gaussian (Wilkinson et al., 1998). Stimulus radius was fixed at 1.5 degrees and the carrier spatial frequency of the narrowband D4 Gaussian was fixed at 2 cycles per degree, which is typically within the range of peak contrast sensitivity for very young animals (Boothe et al., 1988). Stimuli were presented in pseudo-random order.

**Figure 1:**
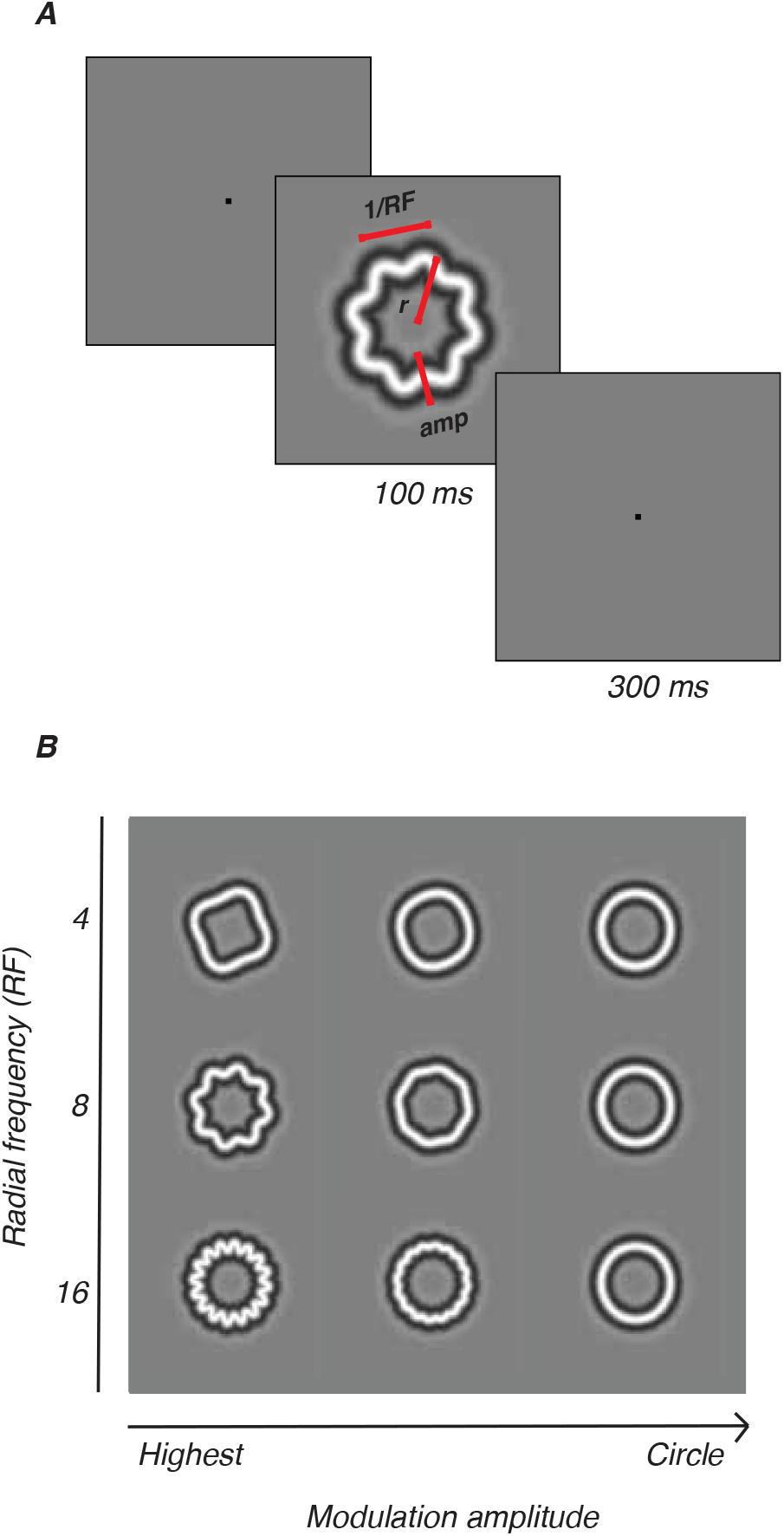
Radial frequency stimuli. A. RFS defined by radial frequency (RF), amplitude (amp), and size (mean radius, r). One trial consisted of 4 interleaved RFS, circles, blanks; 100ms each, 300ms interval. B. Three radial frequencies (4, 8, 16) drawn at various amplitudes.

**Figure 2:**
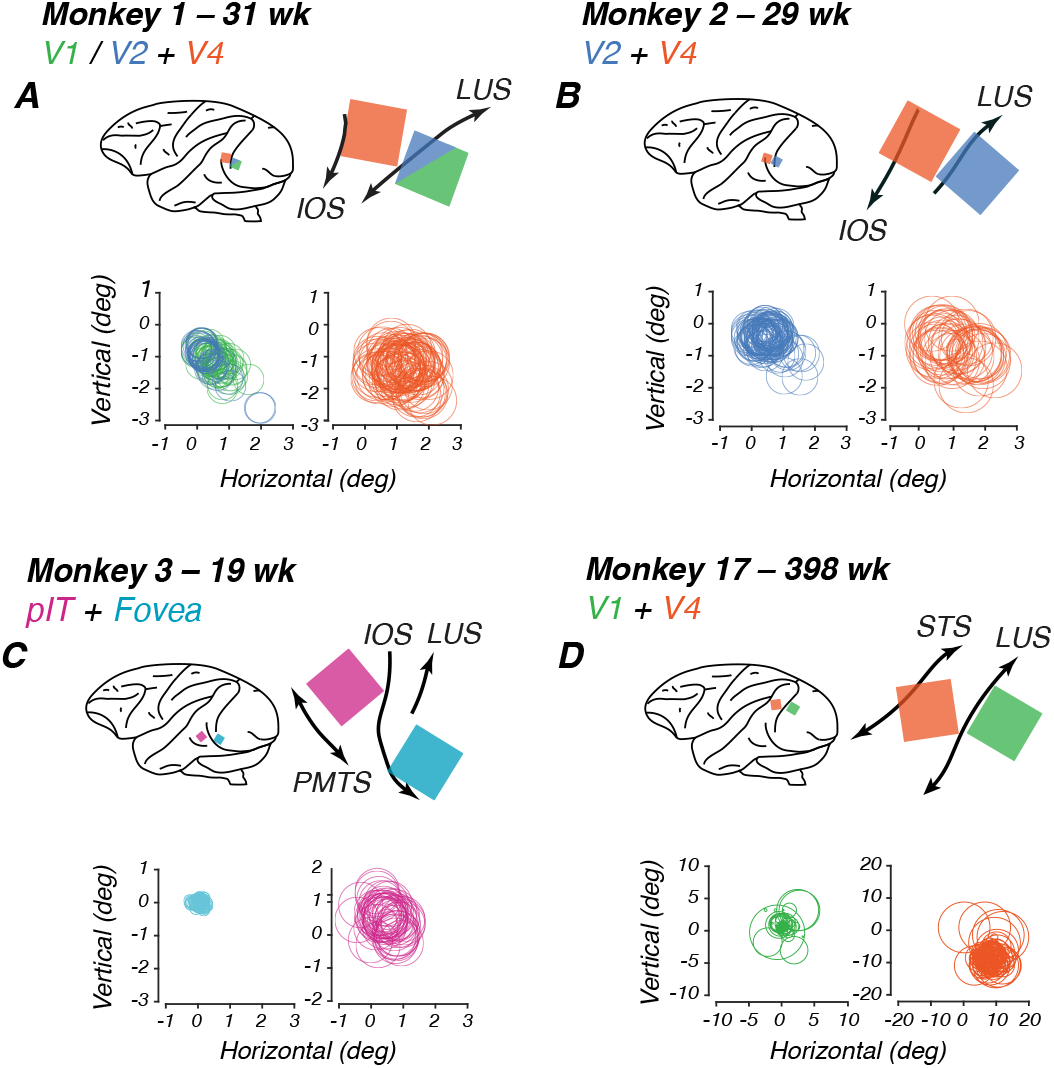
Array sites and receptive field locations for each subject. Each panel shows data for one monkey. In each panel: Top left: schematic lateral view of the brain showing array locations (colored squares) relative to standard anatomical landmarks (black curves). Top right: Detail of Utah array placement according to surgical and histological records. Below: Receptive fields for each array channel. **A**. M1, implanted at 23 wk. **B**. M2, implanted at 23 wk. **C**. M3, implanted at 17 wk. **D**. M17, implanted at 384 wk. Color code: V1 (green), FC (teal), V2 (blue), V4 (orange), pIT (pink). Abbreviations: STS = Superior temporal sulcus; IOS = Inferior occipital sulcus; LUS = Lunate sulcus; ECS = Exterior calcarine sulcus.

### Tasks

Eye tracking signals were calibrated at the beginning of every recording session. The calibration task consisted of a set of trials during which the subject was required to fixate on a 1-degree square that appeared in different locations on the screen on each trial.

#### Receptive Field Localization

Animals fixated on a small dot on the screen while patches of bandpass-filtered white noise were flashed for 100 ms within a 5×5 grid of possible locations around the fixation point in translational steps of 0.25-1°. We fit the threshold crossings for each channel with a Gaussian, with the receptive field center located at the peak of the curve. By shifting each channel such that the maximum of their response grids overlapped, we fit a two-dimensional Gaussian function and used its full width at half maximum as the average size of the population receptive field for each array. Stimuli were presented in the receptive field location identified for each animal. For the infants, receptive fields were roughly foveal and aligned across both arrays, which allowed us to record simultaneously from both arrays. Our adult’s V1 and V4 receptive fields were not as closely aligned and data were collected at separate locations for each array (see Results, Figure 2 and Table 2).

**Table 2:** Mean receptive field diameters and locations, and receptive field diameter-to-stimulus-size ratios for each array.

| Subject | Cortical area | Mean diameter<br>(deg) | Diameter-to-stimulus<br>size ratio | Mean center location<br>(azimuth, elevation) deg |
| --- | --- | --- | --- | --- |
| M1 (female) | V1 | 0.95 | 0.32 | (0.6, -1.1) |
|  | V2 | 0.96 | 0.33 | (0.4, -1.1) |
|  | V4 | 1.52 | 0.51 | (1.2, -1.4) |
| M2 (female) | V2 | 1.14 | 0.38 | (0.6, 0.5) |
|  | V4 | 1.72 | 0.57 | (1.2, -0.8) |
| M3 (male) | FC | 0.37 | 0.12 | (0.1, 0.1) |
|  | pIT | 1.38 | 0.46 | (0.6, 0.5) |
| M17 (male) | V1 | 0.93 | 0.31 | (0.13, 0.68) |
|  | V4 | 4.02 | 1.34 | (7.91, -8.43) |

#### Rapid Serial Visual Presentation (RSVP) paradigm

Animals fixated on a point on the screen for a required amount of time while stimuli flashed over the receptive field of the arrays. The diameter of the stimuli spanned 3 degrees of visual angle in total. An RSVP trial began when the animal was fixating on a red fixation point (0.1-0.2 degrees) presented either at the center of the screen (M1-M3) or adjusted to the upper left, to account for the location of M17’s receptive fields. Following a 160 ms interval, a train of four images appeared, with each image being presented for 100 ms and interstimulus interval of 300 ms (Figure 1A). For a complete trial, subjects were required to maintain fixation for 1760 ms. We tested radial frequencies of 4, 8 and 16 cycles, each with five modulation amplitudes. The modulations were specified as Weber Fractions (WF) - the proportion of radius modulation relative to the base circle radius. The tested modulations were 6.25, 12.5, 25, 50, and 100 (WF × 1000), corresponding to approximately 0.019, 0.038, 0.075, 0.15, and 0.3 degrees of visual angle (deg) for our stimulus size. These stimuli were interleaved with blanks and base circles. If the animal broke fixation prematurely, stimuli disappeared from the screen and the trial was aborted. An error tone would play, followed by a brief time out before the beginning of the next trial. If the animal broke fixation on five consecutive trials, a longer time out would be triggered during which the gray background screen would turn a uniform red. To maintain the level of visual adaptation, the intensity of the red was set to be perceptually isoluminant with the mean gray background.

### Analyses

#### Significance Testing

To determine the probability that a measurement could have occurred by chance, significance testing was done with permutation tests, unless otherwise noted. Stimulus identities were randomized and the corresponding measure was computed on each iteration. We carried out 100-1000 iterations, depending on the computational demands of the analysis. Experimental values greater than 97.5% or less than 2.5% of the permuted distribution were taken to be significant.

#### Post-stimulus time histograms

The data from each recording are first plotted as post-stimulus time histograms (PSTHs) to visualize stimulus-evoked responses on the array (see Results, Figure 3, top row). The mean response to a blank screen was subtracted from the mean stimulus-evoked response to the highest modulation amplitude and plotted as a thick line. Black boxes along the abscissa indicate stimulus duration relative to the response traces. Response latencies were calculated using a half-peak method. For each PSTH, we first identified the peak response value within the defined response window. We then set a threshold at half of this peak value and searched for the first time point where the response crossed this threshold.

**Figure 3:**
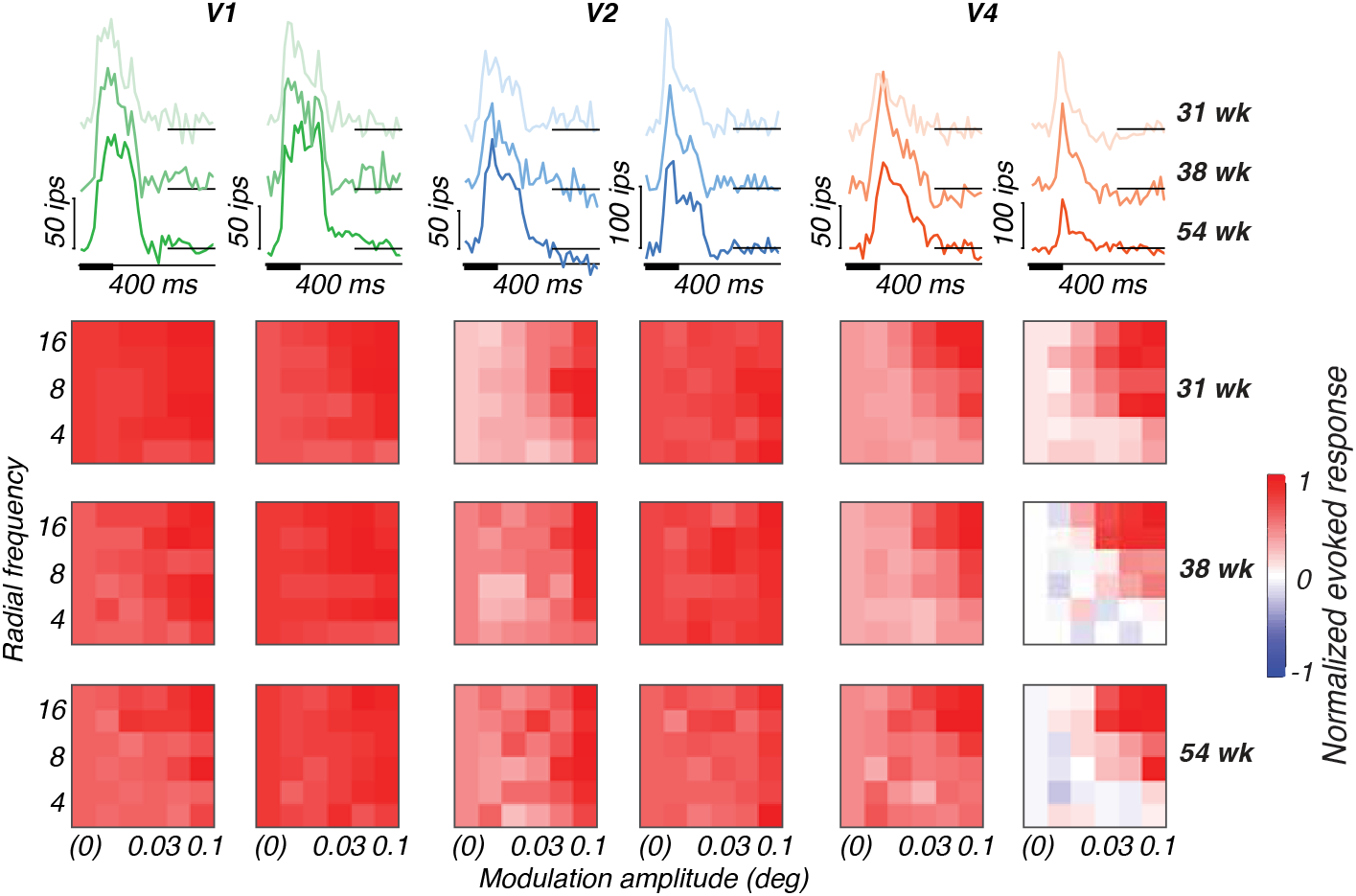
Example response properties of single channels for subject M1. Channel responses to highest modulation amplitude stimuli (top row) at multiple ages (lighter color = younger; darker color = older) and corresponding tuning heat maps (bottom rows) for each animal at multiple ages, with responses plotted as a function of modulation amplitude (abscissa), radial frequency (ordinate) and rotational phase (interleaving rows). Colors represent cortical areas (green = V1; blue = V2; orange = V4).

#### Visual responsiveness

The stimulus vs. blank d’ was used to quantify visual sensitivity for each channel.

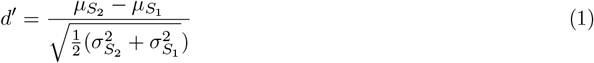

Here, *µ* and *σ* correspond to the mean and variance of the responses to two different stimuli. Responses to a blank screen were denoted as *S*_1_ and responses to the highest modulation amplitude for each radial frequency type were denoted *S*_2_. Each set of neural response data was z-scored across all stimulus conditions. Our responsivity-based exclusion criterion was a *d* less than 0.7, because this ensured the exclusion of channels that were clearly contaminated by noise.

### Tuning variance explained

The percentage of the total tuning variance that could be accounted for by each stimulus dimension (radial frequency, modulation amplitude and phase) was computed by first baseline-subtracting responses across a 50-250 ms interval post-stimulus onset and normalizing by the peak response of each channel.

Resulting responses were visualized by arranging them into 6×6 heat maps with modulation amplitude on the abscissa, radial frequency on the ordinate, with stimulus phase depicted on interleaved rows (e.g., Figure 3, bottom rows). We used these data to compute average tuning vectors and their variance for each stimulus dimension. Tuning variance was normalized by the total variance across the entire heat map to obtain a measure of tuning variance explained by either radial frequency type, modulation amplitude or stimulus phase. Channels that were significantly tuned for at least one stimulus dimension were identified by a permutation test. The 6×6 tuning heat map was randomly permuted and tuning variance explained was computed on each iteration. We repeated this process 100 times and used the distribution of estimated tuning variance explained values to compute the probability that the observed tuning variance explained may have occurred by chance for any of the stimulus dimensions. This was done per stimulus position. The percentage of channels responsive to each stimulus dimension was determined by calculating the proportion of channels that showed significant responsiveness. We then determined the percentage of responsive channels with significant tuning variance explained for radial frequency, modulation amplitude, or stimulus phase.

### Triplot representation

We compared the amount of variance explained by each stimulus dimension using a “triplot” representation similar to that used by (El-Shamayleh and Movshon, 2011). The tuning of each channel was defined by 3 values: the variance explained by radial frequency type, modulation amplitude and rotational phase. These three normalized values define a unit vector in three dimensions, which correspond to a position on the surface of a sphere. Our triplot representations take advantage of the fact that the correlations are always positive to represent the data in one orthant, whose projection onto the plane is shown. The distance between each point from each edge of the triangle is proportional to the amount of tuning variance explained by the labeled dimension (see Figure 4). We represent the amount of variance not explained by the three stimulus variables for that channel by color-shading, with darker points indicating channels where our stimulus dimensions account for most of the response variability, suggesting robust tuning, while lighter points indicate channels where responses are largely driven by other factors or noise. This distinction helps to identify the most reliably tuned channels in each area, beyond just their relative selectivity for different stimulus features.

**Figure 4:**
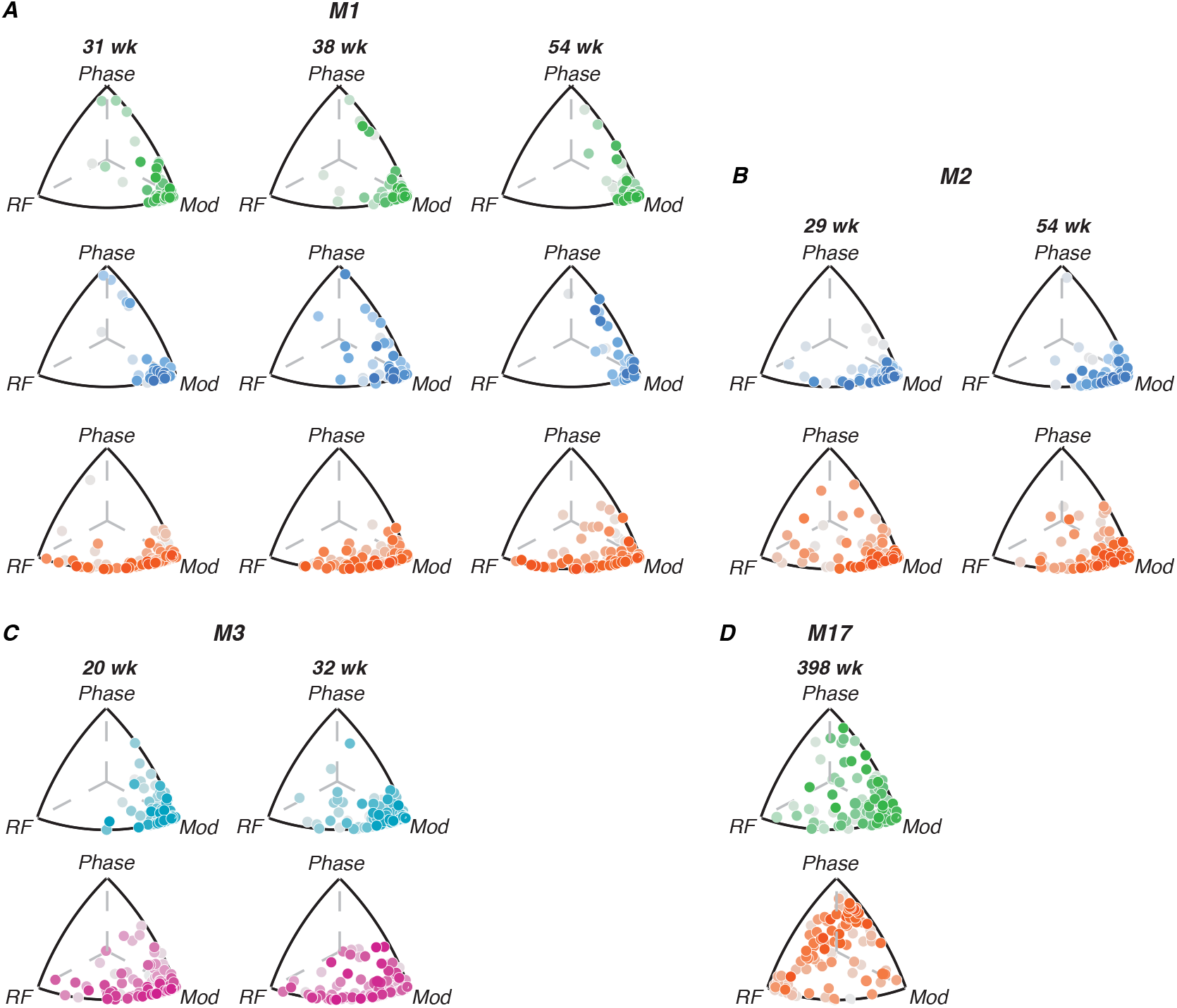
Contributions of radial frequency stimulus features to single-channel tuning. Each point represents a single channel; it’s position relative to each edge of the triplot represents how much variance of the channel’s tuning is explained by radial frequency stimulus variables (radial frequency, modulation or phase). The points are color-coded by the amount of variance not explained by either of the three variables of interest; darker shades represent stronger tuning. Color code: V1 (green), FC (teal), V2 (blue), V4 (orange), pIT (pink).

### Optimal stimulus position

Radial frequency stimuli were presented at 3 locations on the screen according to the receptive field mapping done at the beginning of the experimental timeline. The 3 locations were chosen such that no single channel could cover the entire stimulus, while ensuring the stimulus fell on a large portion of the population receptive field. The optimal stimulus position for analysis of each array was defined as the position that generated the largest amount of tuned and responsive channels. Results presented here correspond to the optimal location for each array.

### Channel inclusion criteria

To compensate for changes in array recording quality across time, we excluded poor-quality recordings based on the number of responsive and tuned channels on the array. Any data set with less than 10 tuned and responsive channels at the optimal stimulus position for each subject was excluded from our analysis. We then combined trials across recordings per day, z-scored daily data sets and ran the d’ and tuning variance explained analyses on this day-by-day data. Days with less than 10 tuned and responsive channels were excluded. Trials for “good” days were subsequently combined and z-scored per week, analyzed for responsiveness and tuning variance and any weeks that failed to meet the aforementioned quality requirements were excluded. Each week is considered an individual age point. This way, we established the minimum number of tuned and responsive channels across ages for each subject. We matched the number of channels across data sets for a given subject by sub-sampling channels from the sets with larger numbers of reliably tuned sites.

### Channel location within extrastriate cortex

We used a combination of gross anatomical features and response properties to confirm which cortical area was being recorded by each channel. Surgical photographs at the time of implantation afforded us the physical layout of the arrays and reference sulci. We refined our estimates of array placements using anatomical atlases and previous reports of electrode locations in adult macaques (Fenstemaker, 1986; Gross et al., 1972; Saleem and Logothetis, 2012; Winters et al., 1969). For M1’s posterior array, located along the V1/V2 border–as confirmed by the visible transition in vascular density on the cortical surface (Zheng et al., 1991)–35 of the 96 channels exhibited slightly slower temporal dynamics, consistent with the response properties of cells in V2.

### Quantifying neural sensitivity

We chose radial frequency stimuli so that a single channel or neuron could not encode the entire stimulus. We used two population decoding methods to characterize the pattern of responses elicited from each population by each radial frequency stimulus. We measured population classification accuracy and created neurometric functions to capture the performance of the population. To calculate population performance, we first established the maximum number of trials and reliably tuned and responsive channels that allowed equating across all recorded age points for each subject. We bootstrapped the data to match these numbers across data sets to ensure cross-validation.

#### Maximum correlation coefficient classifier

We primarily used a correlation coefficient classifier because it models neural computation through synaptic weight integration, making its decoding accuracy biologically plausible (Majaj et al., 2015; Meyers et al., 2008). Training of the correlation-based classifier consisted of creating a classification vector that corresponds to the difference in response between the two stimuli our observers discriminated, radial frequency stimulus and circle. Each classification vector was defined as the difference between the mean response to RFS and the circle, with the length of the vector corresponding to the number of channels. To assess discrimination performance, four trials were drawn, each a vector of evoked neural responses, from our test RFS and circle data sets. Three of the trials corresponded to circle responses and the fourth corresponded to a radial frequency stimulus of a particular modulation amplitude. A Pearson correlation coefficient was calculated between each trial and the classification vector and the trial with the highest correlation was picked as corresponding to an RFS. Those trials were scored and performance was tallied as percent correct. Performance accuracy is reported as the percentage of correctly classified trials. Neurometric functions were generated based on decoding performance accuracy for the range of modulation amplitudes tested for each RFS.

We analyzed the impact of trial-to-trial variability on decoding by training the classifier on data in which trial order was shuffled randomly and independently for each site, to disrupt the correlation structure shared by each population.

#### Linear discriminant analysis (LDA)

Consistent with recent work analyzing texture perception in these same animals (Lee et al., 2024), we also analyzed our neural populations’ ability to discriminate between circles and radial frequency patterns using LDA as a comparison metric. We computed the orientation of the hyperplane that separated the data such that channels whose mean responses were farther apart between stimulus classes would receive larger weight coefficients and were considered most informative for discrimination. We quantified the population’s sensitivity to radial frequency stimuli by comparing the distribution of responses of a population to a circle and a radial frequency stimulus of a given modulation amplitude for each channel and subsequently averaging across channels. We then projected our response data onto that hyperplane and computed the d’ metric of discriminability for the population response.

Both methods yielded qualitatively similar results, demonstrating the robustness of our findings across different decoding approaches.

## RESULTS

To measure neural sensitivity to global form stimuli, we implanted two 96-channel “Utah” arrays in each animal. We trained the animals to maintain fixation on a dot while a sequence of stimuli was flashed in random order. We only analyzed channels that passed our inclusion criteria of being visually responsive and significantly tuned to any stimulus dimension, unless otherwise specified. Array placements are shown below in Figure 2.

### Receptive field localization

To locate where the receptive fields were concentrated for each multi-electrode array, we fit threshold crossings in response to filtered noise stimuli and plotted the center of the receptive field as the peak of the fitted Gaussian for each channel. Circles enclosing the receptive fields for each channel are plotted in Figure 2, yielding composite receptive fields for the population recorded by each array. In each panel, the fixation point lies at the origin of the axes, and every circle indicates the estimated location of one channel’s receptive field. Measures of these receptive field estimates are listed in Table 2. For our infants M1-M3, population receptive fields in V1/V2, V4 and pIT were largely overlapping and foveal, allowing us to evoke activity in both arrays using the same stimulus locations in the central visual field and compare to our behavioral results. For subjects M1-M3, arrays yielded average receptive field sizes and centers located near the center of gaze. For our youngest subject, M3, the receptive field sizes and response properties suggest that his posterior array was located at the foveal confluence (FC) of areas V1 and V2. For our adult subject, M17, V1 receptive fields were mostly within 2 deg of the fovea. However, his V4 receptive fields were larger and centered roughly 12 deg in the periphery. The goal of our mapping routine was to identify the general locations that our arrays were responsive to, so that we could optimize the location of the radial frequency stimuli on the screen to ensure that the population receptive field was covered by a significant portion of the stimulus. To quantify this relationship, we calculated the ratio between receptive field diameter and stimulus diameter (3 deg) for each array (Table 2). This ratio measures how much of the stimulus could fall within a single receptive field, with values greater than 1 indicating receptive fields large enough to encompass the entire stimulus.

### Response properties

#### Single-site responsiveness across age

To characterize the response properties recorded on each multi-unit channel of the arrays, we plotted baseline-subtracted PSTHs at the best stimulus location. In Figure 3 (top row), we show two example channels per array for subject M1. In the top row for each subject, the mean baseline-subtracted response (imp/s) to the highest modulation amplitude stimuli across radial frequency is plotted as a function of time. Each trace corresponds to responses at different ages, with color saturation increasing across developmental time. Stimuli were presented for 100 ms (thick bar on abscissa) and remained off for 300 ms after each presentation. As reflected by the example channels presented here, we observed a range of response profiles across ages and arrays. The range of firing rates across channels on our various arrays were typically between 20-300 imp/s, with channels most commonly responding in the 20-150 imp/s range. Notably, we did not observe systematic changes in tuning profiles, response shape or latency as a function of age (Table 3). However, the typical progression in response latency across cortical areas is observable at all ages studied.

**Table 3:** Response latency (*mean ± S*.*D*.) by age and area.

| Cortical area | Age group (wk) | Latency (ms) |
| --- | --- | --- |
| V1 | 30-35 | 58.4 ± 9.5 |
| | 40-45 | 57.4 ± 4.6 |
| | 50-55 | 53.4 ± 12.3 |
| | > 380 | 59.4 ± 22.1 |
| V2 | 30-35 | 70.4 ± 11.0 |
| | 40-45 | 70.0 ± 11.1 |
| | 50-55 | 66.8 ± 16.6 |
| V4 | 30-35 | 73.0 ± 32.2 |
| | 40-45 | 81.0 ± 23.4 |
| | 50-55 | 67.9 ± 27.4 |
| | > 380 | 70.3 ± 13.5 |
| pIT | 19-22 | 85.8 ± 50.6 |
| | 25-33 | 83.2 ± 52.9 |
| FC | 19-22 | 74.8 ± 43.9 |
| | 25-33 | 62.0 ± 35.9 |

To address how responsive channels compared across stimulus types and ages, Table 4 shows the percentage of channels responsive to both filtered noise stimuli (used for receptive field mapping) and radial frequency patterns, organized by cortical area and overlapping age groups. Overall, we observed substantial responsiveness to both stimulus types across all areas and ages, with some variability that did not show a consistent developmental pattern for either stimulus type. V1 showed particularly high responsiveness to both stimulus types. Areas like V2 and V4 demonstrated greater relative responsiveness to radial frequency patterns compared to filtered noise in some age groups, but not others. FC and pIT showed relatively higher responsiveness for filtered noise stimuli. It is important to note that several factors may influence these percentages, including stimulus-specific properties (filtered noise patches versus contour-based radial frequency patterns) and technical considerations such as subject-to-subject variations in array recording quality. However, these data complement our PSTH analysis, showing fairly stable response properties throughout developmental time.

**Table 4:** Percentage of channels responsive to filtered noise and radial frequency stimuli by age group and area.

| Cortical area | Age group (wk) | Radial frequency (%) | Filtered noise (%) |
| --- | --- | --- | --- |
| V1 | 30-35 | 97 | 89.6 |
|  | 50-55 | 96.9 | 75.6 |
|  | > 380 | 59.4 | 65.5 |
| V2 | 30-35 | 70.5 | 43.3 |
|  | 40-45 | 77.6 | 33.9 |
|  | 50-55 | 63.5 | 64.5 |
| V4 | 30-35 | 34.2 | 17.6 |
|  | 40-45 | 37.2 | 2.9 |
|  | 50-55 | 27.1 | 68.2 |
|  | > 380 | 95.8 | 63.5 |
| pIT | 25-33 | 9.8 | 61.6 |
| FC | 25-33 | 20.8 | 58 |

**Table 5:** Tuning to radial frequency pattern features across cortical areas. The percentage of responsive channels are reported. Columns show the percentage of responsive channels tuned to different stimulus features. Significant values are denoted in bold font.

| Cortical area | Responsive (%) | Frequency-tuned (%) | Mod-tuned (%) | Phase-tuned (%) |
| --- | --- | --- | --- | --- |
| V1 | **88.6** | 9.8 | 40.2 | 8.9 |
| V2 | **66.9** | 12.9 | 59.2 | 9.3 |
| V4 | 31.1 | **26.7** | 48.6 | 7.8 |
| pIT | 21.4 | 17.3 | 34.7 | 5.9 |
| FC | 8.4 | 3.3 | 37.7 | 4.4 |

#### Tuning to radial frequency stimuli

To visualize the stimulus selectivity profiles of each channel, we plotted the baseline-subtracted firing rate normalized to its peak response as tuning heat maps, in which the intensity of the color represents the magnitude of the response for a given condition (Figure 3, bottom rows). The patterning of excitatory (red) and suppressive (blue) activity across stimulus conditions reflects varying degrees of selectivity to global form features on a channel-by-channel basis across the array. Each heat map shows how an example channel responded based on modulation amplitude (abscissa), radial frequency (ordinate) and rotational phase (interleaved rows). Beyond exhibiting selectivity for radial frequency, single channels displayed differential responses to particular stimulus features such as phase and amplitude modulation. How these stimulus features were processed depended on both cortical area and receptive field size relative to the stimulus. For example, channels in V1 were broadly responsive across stimulus conditions. In V2 and to some extent FC, some channels exhibited selectivity for the larger modulations across radial frequency patterns. In V4, we observed channels with a range of selectivity profiles. Some channels were narrowly-responsive to the highest modulation and radial frequency pattern, for example. We also observed narrowly-tuned channels in pIT (not shown), but not as many as we observed in V4. Notably, some of M17’s V4 channels (not shown here, but see Figure 4D) were remarkably selective for stimulus rotation (phase), reflected by horizontal bands in their tuning. This distinctive phase selectivity may reflect a difference in visual processing at different spatial scales. M17’s peripheral V4 had receptive fields large enough to encompass the entire 3° stimulus (receptive field/stimulus ratio = 1.34 (Table 2). In contrast, the more foveal V4 receptive fields in younger animals had ratios of 0.51-0.57, suggesting their receptive fields could only process portions of the stimulus at a time. The larger receptive field/stimulus ratio in M17’s V4 may have enabled the neurons contributing to these channels to encode rotational phase. This interpretation suggests that receptive field/stimulus size ratio may be a key determinant of a neuron’s ability to encode global stimulus features like phase as local oriented fragments or as global stimulus rotation.

Figure 4 illustrates the relative contributions of different radial frequency stimulus dimensions to single-channel tuning across visual areas and developmental time for each animal. Each point represents a single channel, with its position in the triplot reflecting the proportion of tuning variance explained by radial frequency, modulation amplitude, and rotational phase. The color intensity indicates the percentage of noise variance, with darker points representing stronger tuning. In early visual areas (FC/V1/V2), tuning is primarily driven by modulation amplitude across all ages, as evidenced by the clustering of points near the “modulation” vertex. This suggests that neurons in these areas are most sensitive to local contrast changes rather than global form features. In area V4, neurons show a mixture of selectivity for radial frequency and modulation amplitude, indicating an emerging selectivity to global form in this region. For M17, V4 neurons exhibit strong phase tuning. In pIT, tuning appears to be driven mainly by radial frequency and modulation amplitude but not to the same extent as V4. Comparing across areas, we see a clear progression in the complexity of tuning from early visual areas to V4, with pIT potentially inheriting a degree of selectivity from V4 that contributes to its role in high-level object recognition.

Importantly, V4 neurons show some degree of global form sensitivity even at the earliest ages we tested.

As illustrated in Figures 3-4 and Table 4, we did not observe systematic changes in tuning across the ages sampled for M1-M3, but did observe inter-areal differences in tuning profiles. To quantify this, we aggregated data from all channels across animals and ages, grouping by cortical area. We then computed both the percentage of channels responsive to radial frequency stimuli and, among responsive channels, the proportion tuned to each stimulus dimension. The second column in Table 4 shows the weighted-mean percentage of responsive channels on each array, compiled across all channels, animals and arrays. A Kruskal-Wallis test revealed significant differences between areas in the percentage of responsive channels (*H*(4) = 31.02, *p <* 0.01, *ε*^2^ = 0.74), with V1 and V2 yielding significantly more active channels across subjects than FC and pIT (Dunn’s test; pairwise comparisons *p <* 0.05). V1 also had significantly more active channels than V4 (*p <* 0.01), while V2 and V4 did not differ significantly.

The following three columns show the percentage of responsive channels tuned to each stimulus dimension of interest. We found significant differences between areas in radial-frequency-selective channels (*H*(4) = 22.67, *p <* 0.01, *ε*^2^ = 0.54), with V4 having more channels selective to this dimension than V1 (*p <* 0.01) and FC (*p <* 0.001). pIT shows a trend toward having more radial frequency-tuned channels than FC, though this doesn’t reach statistical significance (*p* = 0.06). We did not find significant differences between V1 and V2, V2 and V4, or V4 and pIT in terms of radial frequency tuning. V2 was particularly well-tuned to modulation amplitude, consistent with its known role in extracting local statistical content of visual stimuli. However, this difference was not statistically significant from other areas. All arrays in the infants exhibited little tuning to rotational phase. The exception to this trend was the adult - M17’s - V4, which exhibited a particularly interesting pattern of selectivity to rotational phase. The relative proportions of channels tuned to each stimulus dimension were largely consistent with our expectations given the receptive field sizes measured in each cortical area, as noted above. These results support the hierarchical processing model of global form information, with V4 emerging as particularly important for radial frequency pattern encoding compared to earlier visual areas like V1. The increasing sensitivity to radial frequency moving from V1 to V4 suggests progressive integration of local orientation information into global shape representations along the ventral stream.

### Population decoding

Our analysis of single-site tuning to global form features in extrastriate regions showed clear RFS tuning even at the earliest ages tested in this study. Two factors make it unlikely that single neurons could support an observer’s ability to discriminate between circles and other shapes: First, individual neurons show considerable trial-to-trial response variability that limits their reliability (Tolhurst et al., 1983).

Second, the synaptic architecture of the visual cortex necessitates population-level processing rather than direct readout from single neurons (Britten et al., 1996; Shadlen et al., 1996). Our choice of radial frequency stimuli necessitates studying population responses since a single channel in central vision cannot encode the entire stimulus form. A population of neurons is needed to signal the global neural response to these stimuli. To address this, we used a population decoding analysis across the ventral stream at different developmental time points.

### Neural sensitivity to global form stimuli

To evaluate how neural populations might support task performance, we implemented a maximum correlation coefficient classifier to simulate our animals’ 4-choice task. This classifier compares test patterns to templates based on response shape rather than absolute firing rates, making it robust to firing rate fluctuations - an important factor for our analysis of developmental data. For each subject, channels that were tuned for at least one stimulus dimension at the optimal stimulus location for the array were included in our population decoding analyses, establishing the largest common set of tuned channels across developmental time and cortical areas for each subject.

In Figure 5, we present examples of population decoding performance for all four animals. Population classification accuracy is plotted as the proportion of correct trials for each radial frequency stimulus, generating neurometric functions describing the neural population’s performance. Each row depicts data for a given subject at the earliest age tested for each, with performance per radial frequency in each column. The curves in each panel represent population decoding performance, with each color representing the cortical area recorded (see figure legend). From this figure, we can appreciate that V4 usually outperformed all other areas. V2 showed good performance, especially at high radial frequency. FC and V1 gave no reliable signals for this task, while pIT showed some sensitivity, but less than V4 or

**Figure 5:**
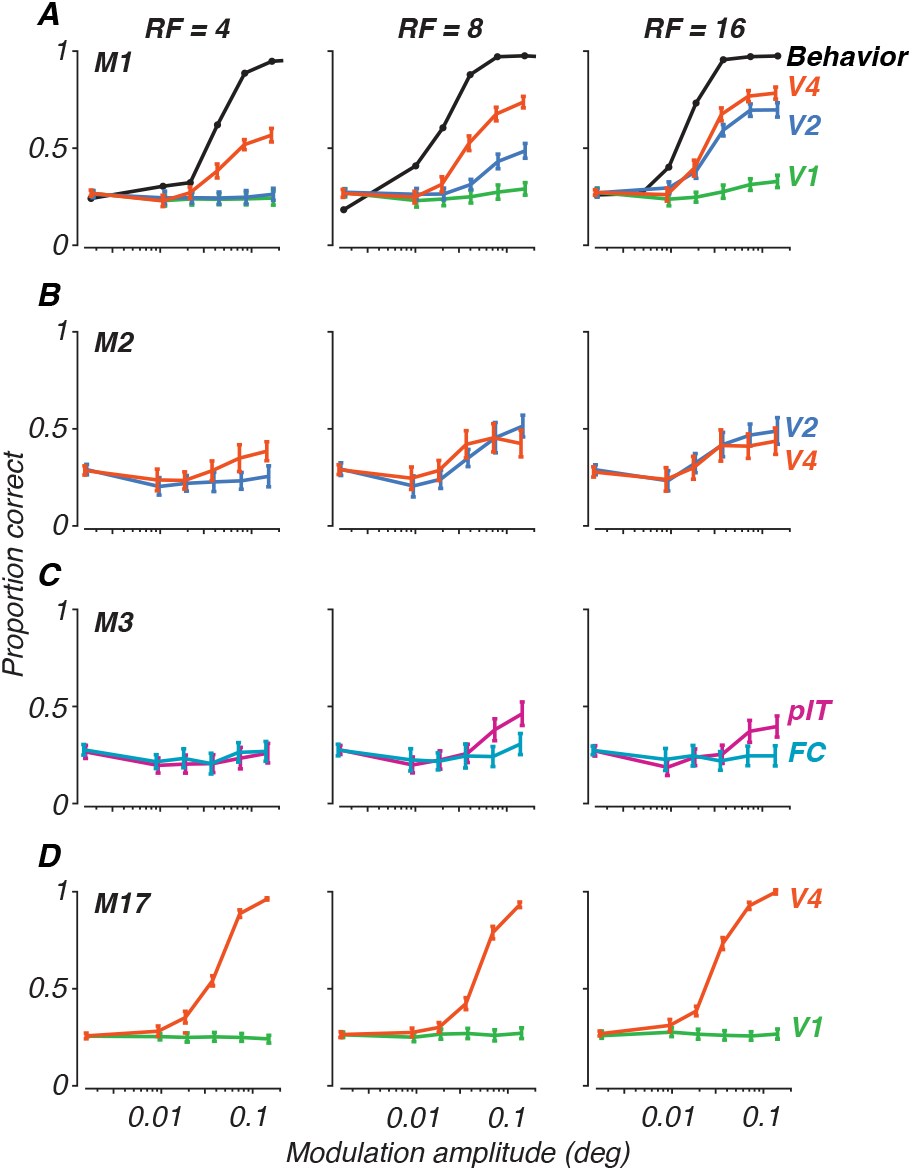
Population decoding of radial frequency stimuli on a 4-AFC task. Each panel plots discrimination performance against modulation amplitudes for each radial frequency type (columns). Error bars represent standard deviation across trial bootstraps. Each row represents a subject. **A**. Monkey M1, age 31 wk, decoded from 32 sites. **B**. Monkey M2, 29 wk, 28 sites. **C**. Monkey M3, 19 wk, 11 sites. **D**. Monkey M17, 398 wk, 57 sites. Colors represent cortical areas (green = V1; teal = FC; blue = V2; orange = V4; pink = pIT).

V2. These patterns were consistent across both our correlation coefficient classifier and LDA approaches, supporting the robustness of these findings and consistent with our work on texture sensitivity (Lee et al., 2024). For comparison to behavior, we have added psychometric functions measured at 17 weeks for one animal, M1 (Figure 5, panel A), on the same scale, illustrating the differences in overall performance between neural decoding and psychophysics and modulation sensitivity for the behavioral and neural data (Rodriguez Deliz et al., 2024). Inspection of the neurometric functions in Figure 5 illustrates why it would be difficult to quantitatively measure thresholds in the case of the neural data: many of the curves do not reach high performance levels even at the highest modulation. Direct comparison between neural and behavioral thresholds is therefore not possible.

We conducted our population classification analysis using two approaches to evaluate the role of local versus global cues in neural discrimination. First, we trained and tested the correlation classifier using data from both stimulus phases combined, which allowed us to assess overall discriminability regardless of stimulus orientation - represented here as phase. We then compared these results to a more stringent analysis where we trained the classifier on responses collected for one stimulus phase and tested it on the second phase. This cross-phase validation approach was designed to ensure that orientation-specific local cues present in the stimulus were not driving the classifier’s decisions. Importantly, we found no substantial differences between these two approaches across any of the visual areas studied, including M17’s V4 where strong phase selectivity was observed. The lack of difference between phase-combined and cross-phase analyses suggests two key points: first, that phase was not processed as a local cue even in large receptive fields capable of encompassing the entire stimulus, and second, that phase selectivity as a global feature did not affect discrimination between radial frequency patterns at different orientations. While other local cues beyond phase might be present in our stimuli, the consistency between phase-combined and cross-phase analyses suggests that neural populations were encoding stable stimulus features rather than phase-dependent local cues, whether local or global. Given the consistency of results across methods, we present the cross-phase validation results in Figures 5 and 6. V1 and FC showed consistently low discrimination performance in both analyses, while V2, V4, and pIT maintained similar levels of discrimination performance regardless of whether phases were combined or separated in the training and testing sets. Our results demonstrate that phase-invariant processing is present across the visual hierarchy, even at the earliest ages tested.

**Figure 6:**
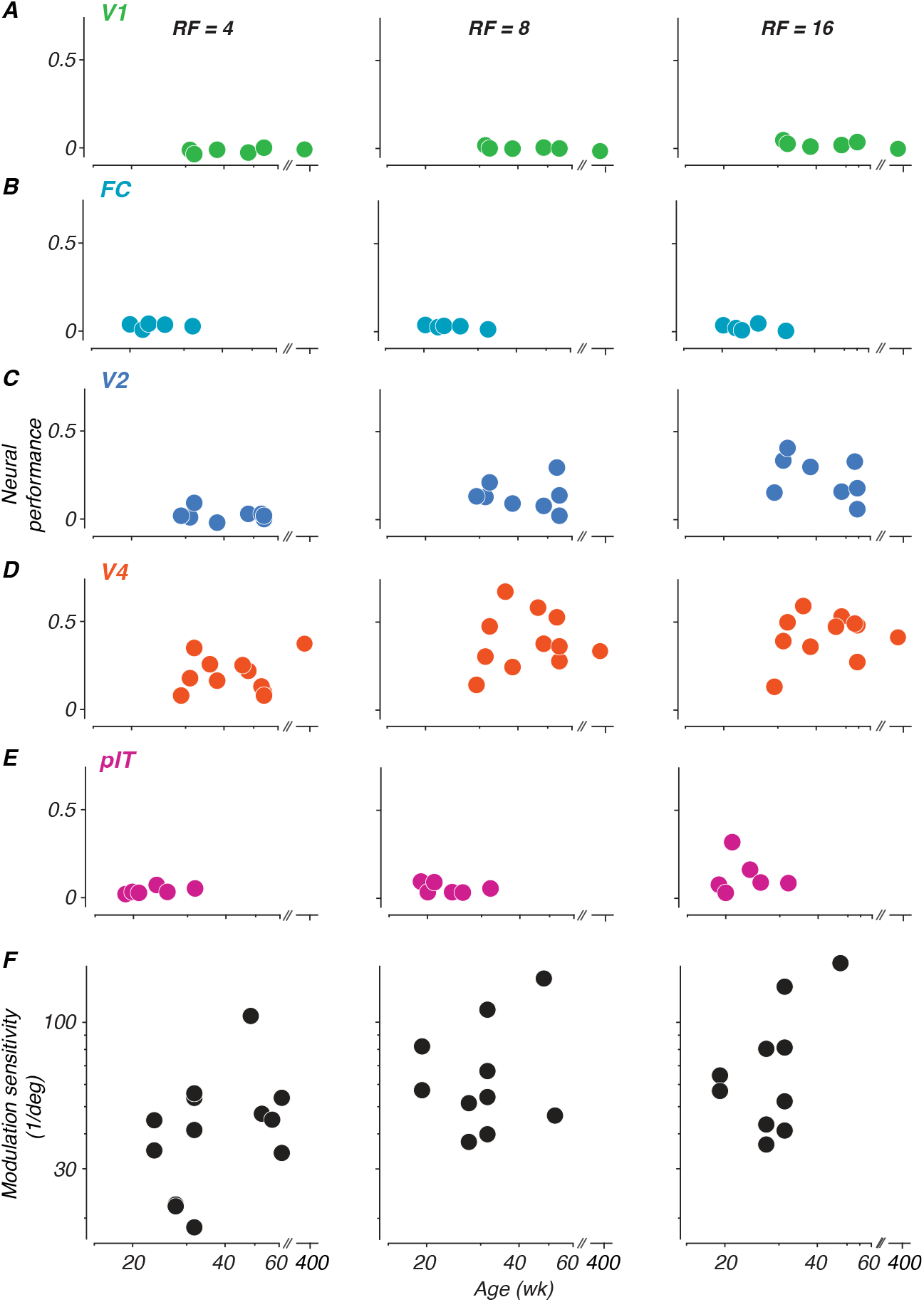
Development of neural sensitivity to radial frequency stimuli. (A-E) Each panel depicts neural sensitivity as summarized by the normalized sum of the neurometric curve for each age at which neurophysiological data were recorded for each subject. Columns represent radial frequencies of 4, 8, and 16. Colors represent cortical areas (green = V1; teal = FC; blue = V2; orange = V4; pink = pIT). (F) Perceptual sensitivity to radial frequency stimuli measured with 4-choice oddity discrimination task. Subset of data excerpted from Figure 5 of (Rodriguez Deliz et al., 2024). All subjects reported in this study were part of the cohort tested on the behavioral task.

Superimposed on the data for monkey M1 in **A** are behavioral psychometric functions for the same animal at the age of 17 weeks (from Rodriguez Deliz et al. (2024)).

### Tracking neural sensitivity to global form across development

To measure changes in neural sensitivity across age, neural performance was summarized by subtracting the proportion correct at chance level from the proportion correct at each modulation amplitude level, and summing across all modulations. This sum was divided by the number of modulation levels to normalize across different sampling densities, and then divided by 0.75 (the maximum possible difference between perfect performance at 1.0 and chance level at 0.25) to scale the metric between 0 and 1. This normalized metric allows direct comparison of neural sensitivity across different experimental conditions and age points. Each panel in Figure 6 shows neural performance values computed for each age point collected for each animal, plotted as a function of age.

What stands out from these results is that neural sensitivity to global form was best in V4, somewhat lower in V2 and pIT, and basically absent in V1 and the foveal confluence. Moreover, sensitivity was better for higher radial frequencies. We tested the data in Figure 6 for significance using a permuted analysis of variance (Anderson, 2001). As one would suppose from inspection of the data, this revealed strong effects of both area and radial frequency on performance (*F* (4, 101) = 23.44 and *F* (2, 101) = 6.68, *p <* 0.001), but no effect of age (*F* = 96.92, *p >* 0.97). We conclude that there is no evidence in our data for an age-related improvement in the neural representation of our global form stimuli in any of the areas we studied.

In a study parallel to this one, we measured monkeys’ sensitivity to global form stimuli using a 4-choice oddity task, in which our animals (including M1-M3 and M17) discriminated between radial frequency patterns and circles (Rodriguez Deliz et al., 2024). Here, we pose the same question in the context of neural decoding. It is challenging to compare behavioral and neural discrimination performance across development, because as seen in Figure 5, our neurometric measures in many cases did not achieve the level needed to compute thresholds for comparison. Instead, to give a qualitative impression of the relationship, we include in panel F behavioral data from an age range selected to match what was studied here, taken from animals tested with a 4-choice behavioral task (Rodriguez Deliz et al., 2024). These data show a roughly two-fold improvement in behavioral sensitivity over the age range represented by the data in panels A-E.

### Impact of correlated variability between neurons on population decoding

Our classifier was trained on the full recorded data set, and therefore had access to its covariance structure. It is therefore possible that changes in the shared neural variability might have affected our conclusions about neural discrimination performance differently at different ages or in different visual areas. We therefore did separate analyses in which we trained a comparison classifier with the covariance structure disrupted by trial shuffling. We found generally modest effects of correlated variability across cortical areas. In areas V2, V4 and pIT the analysis was inconclusive, with some populations seeming to benefit slightly from correlation structure while others showed negligible effects. The effect of correlated variability also did not vary with age, suggesting that neural population interactions were established before the age at which our measurements began. However, we only offer this as a preliminary finding because the brief trial durations in our experiments limited the sensitivity of our analysis. We will report in a subsequent paper the results of a separate analysis of correlated variability in these animals.

## DISCUSSION

In this study, we investigated the development of neural mechanisms supporting global form processing by recording from areas V1, V2, V4 and pIT of infant macaques using radial frequency stimuli. Our goal was to identify a neural correlate of documented behavioral development within the young visual cortex. Our key finding reveals that neurons in area V4 of infant macaques can reliably encode global form information from radial frequency stimuli as early as we were able to record - 30 weeks of age, with little evidence of systematic improvement with subsequent development. The extrastriate ventral stream demonstrates surprisingly mature processing capabilities early in development, through at least the level of V4.

### Neural sensitivity to global form

Our results demonstrate a clear hierarchical progression in the processing of global form information across the ventral visual stream. Neurons in the foveal confluence and V1 proper were visually responsive at the youngest ages but had limited ability to discriminate radial frequency patterns from circles. V2 outperformed V1 in some cases, as did 19-week-old neurons in pIT for the highest radial frequency tested, but neither V2 nor pIT reached the performance levels exhibited by neural populations in V4 even at 30 weeks. Our findings align with previous work positioning V4 as an intermediate shape processing area exhibiting an emergent selectivity for curvature and shape information that is not observed in earlier visual areas (Gallant et al., 1993, 1996; Pasupathy and Connor, 2002; Poirier and Wilson, 2006; Wilson et al., 1997). Importantly, we demonstrate that this specialization of V4 for global form processing is present in early infancy. This suggests that even in young animals, V4 neurons can effectively integrate multiple local curvature signals to construct global shape representations.

A particularly striking aspect of our results was that the increased perceptual sensitivity observed for higher radial frequencies in both humans and monkeys was reflected in the patterns of stimulus-evoked neural activity, both at single-channel and population levels. Higher radial frequencies contain more cycles of modulation along their contour, providing more local orientation and curvature cues. Our data support the idea that V2 neurons are sensitive to the local orientation information falling upon their receptive fields, with higher responsiveness expected based on the larger number of cycles serving as input drive. V4 neurons, with their larger receptive fields, can then integrate local curvature information from V2, benefiting from the larger amounts of local orientation and curvature content in higher radial frequency stimuli. The higher sensitivities for RF8 and RF16 we observed at the perceptual and neural population level may reflect cortical mechanisms dedicated to enhancing hyperacuity signals in V4 that are available fairly early on in life (Brown, 1997; Kiorpes, 1992, 2015; Norcia et al., 1988; Shimojo et al., 1984; Shimojo and Held, 1987; Skoczenski and Norcia, 2002; Wang et al., 2009; Zanker et al., 1992).

Ecologically, this makes sense because more information about the visual environment is available in finer details represented by higher spatial frequencies, as evidenced by analysis of natural images and facial recognition experiments (Bex et al., 2009; Field, 1987).

Our neurophysiological recordings from V4 and pIT provide new insight into how the visual system might resolve this debate between local and global processing mechanisms. V4 neurons showed clear selectivity for RF patterns even at early ages, but importantly, the individual V4 receptive fields in our infant subjects were smaller than the full stimulus (receptive field/stimulus ratios of 0.51-0.57). Despite this, V4 populations reliably encoded global form information, suggesting that local orientation and curvature signals must be integrated across space by downstream neurons to create a complete shape representation. This population-level encoding may be particularly important for processing higher RFs, where the increased circular contour frequency (Jeffrey et al., 2002) provides more local orientation and curvature cues per degree of contour, consistent with perceptual findings of enhanced sensitivity to higher RFs (Rodriguez Deliz et al., 2024). Distributed processing mechanisms in V4 could support both local orientation analysis and global form perception through population coding (Pasupathy and Connor, 2002), helping to reconcile the longstanding debate about local versus global processing of RF patterns. While V4 clearly contributes to this processing, the limited size of its receptive fields implies the need for a higher-level integrator of this distributed shape information. Identifying where and how this population-level V4 information gets transformed into more abstract shape representations remains an important open question. Further investigation of how neurons in pIT encode RF patterns across development could help reveal how distributed local information is combined into coherent global percepts.

### Linking neurophysiology and behavior

In a parallel study, we showed that behavioral performance on global form tasks continued to improve substantially beyond the first year of development (Rodriguez Deliz et al., 2024). Our physiological findings echo a persistent puzzle: despite clear behavioral improvements, we found little evidence for systematic changes in V2, V4 or pIT over the ages we studied. This dissociation between neural and behavioral development mirrors earlier work in the retina, visual thalamus, V1 and V2 (Kiorpes and Bassin, 2003; Kiorpes and Movshon, 2004; Lee et al., 2024; Maruko et al., 2008; Movshon et al., 2005), and extends it to higher-order visual areas that we hypothesized might show developmental changes that could account for behavioral improvement. Several factors might explain this dissociation. Our recordings sampled relatively small neural populations, while behavior likely depends on much larger neural ensembles. Previous population scaling analyses showed improving performance with increased population size, consistent with prior studies (Averbeck et al., 2006; Shadlen et al., 1996; Shooner et al., 2015) and we may be underestimating the full computational capacity of V4.

Alternatively, behavioral improvements might reflect changes in how downstream areas read out and utilize V4 signals, rather than changes in V4 processing itself. The refinement of functional connectivity between V4 and higher-order areas, or within V4 itself, might continue to improve even after basic processing in V4 neurons is established, leading to enhanced perceptual performance (Bourne et al., 2024). The presence of correlated noise between cells in a population could limit the amount of information available for downstream readout of shape representations (Abbott and Dayan, 1999; Averbeck et al., 2006). In this study, we explored this possibility by testing our classifier on trial shuffled data, with results suggesting a limited effect of noise correlations on decoding performance. Future work should examine how correlated variability in V4 changes developmentally (Rodríguez Deliz, 2023),using stimuli that directly engage V4’s shape processing capabilities and comparing noise correlations in foveal versus peripheral populations to understand differences in shape coding mechanisms across the visual field, as suggested by the differences in phase selectivity we observed between infants and adults.

### Methodological considerations

Several technical aspects of our study warrant consideration when interpreting these results, although our core findings remain robust. The challenges of chronic recordings in developing animals meant that we could only follow individual subjects over limited time windows, and array performance inevitably varied over time. Our rigorous inclusion criteria and matching procedures help mitigate these issues.

These criteria were essential for ensuring data quality and helped control for changes in array performance over time. Moreover, temporal analysis of data from the same subjects revealed that neural populations could reliably discriminate between stimuli even within brief time windows, with consistent response timing and discrimination performance across age groups, suggesting that temporal processing dynamics were not a limiting factor. The infants were head-restrained using custom side panels rather than a traditional headpost, potentially introducing additional response variability compared to adult recordings. However, our consistent findings across subjects and sessions suggest this was not a limiting factor.

Importantly, we observed no systematic changes in task engagement or motivation across ages that might have obscured potential improvements in neural performance at older ages. This is further supported by consistent and reliable behavioral performance during interleaved testing sessions throughout development (Rodriguez Deliz et al., 2024). Lastly, it is possible that training on the behavioral task may have induced experience-dependent plasticity underlying enhanced sensitivity to radial frequency patterns. However, we did not find significant changes in neural decoding reflective of the protracted perceptual development observed over the same age range. Furthermore, M17 had no experience with radial frequency patterns prior to these physiology experiments. This suggests that training effects, if any, might not have had a major impact on our overall conclusions.

### Conclusions

This study demonstrates that V4 neurons in infant macaques are capable of representing global form information and supporting shape discrimination at a young age (30 weeks). The fact that neural sensitivity did not show the same prolonged development as behavioral performance suggests that the period over which shape selectivity develops occurs earlier, and/or that our measurements of small neural populations do not fully capture the selectivity that supports behavior.

The early emergence of form processing in V4, particularly for fine-scale deformations, provides novel insights into the developmental primitives of high-level vision. Further work is needed to elucidate the precise temporal unfolding of these neural mechanisms and their relationship to perceptual abilities across a longer developmental trajectory. Examining responses in larger populations and at a wider range of spatial scales will be crucial for understanding the neural basis of form perception and its development.

## Acknowledgements

We are grateful for the contributions of members of the Visual Neuroscience Laboratory to this work, particularly Jessica Fletcher, Tiffany Tang, Kahlia Gronthos, Sophia Francisco-Simón and Michelle Hernandez for training and testing our subjects. Thanks to Tim Oleskiw for providing important insights about analyzing data from V4. Thanks also to the staff of the NYU Office of Veterinary Resources, our subjects, and the Washington National Primate Center. This work was supported by grants from the National Institutes of Health: R01EY024914 (LK, JAM), R01EY031446 (NJM, LK), R01EY005864 (LK), T32MH019524 (LK), F31EY026791 (BNB), F31EY031249 (GML), and F31EY031592 (CLRD).

## Author contributions

N.J.M., L.K., and J.A.M. designed the experiment. C.L.R.D., B.N.B., and G.M.L. conducted the experiments. C.L.R.D. analyzed the data. C.L.R.D, L.K., and J.A.M. wrote the paper.

